# An explainable boosting machine model for identifying artifacts caused by formalin-fixed paraffin embedding

**DOI:** 10.64898/2026.03.10.710815

**Authors:** Valentina Grether, Zoe R. Goldstein, Jennifer M. Shelton, Timothy R. Chu, William F. Hooper, Heather Geiger, André Corvelo, Rachel Martini, Melissa B. Davis, Nicolas Robine, Will Liao

## Abstract

**Background:** Formalin-fixed paraffin-embedding (FFPE) is a widely used, cost-effective method for long-term storage of clinical samples. However, fixation is known to introduce damage to nucleic acids that can present as artifactual bases in sequencing otherwise absent from higher fidelity storage methods such as fresh freezing (FF). Various machine learning methods exist for filtering these variant artifacts, but benchmarking performance can be difficult without reliable truth sets. In this study, we employ a collection of 90 paired fresh-frozen and formalin-fixed paraffin embedded samples from the same tumor to robustly define real and FFPE-derived, artifactual variation and enable objective evaluation of filtering methods. To address existing shortcomings, we propose a novel explainable boosting machine (EBM) model that improves performance, can be easily updated with new data, requires modest computational resources, and is analysis pipeline agnostic, making it broadly accessible.

**Results:** We evaluated several methods for limiting FFPE-derived variant artifacts using cohorts of B-cell lymphoma samples. We found capturing local context around variants to be a highly informative, under-utilized feature set not commonly incorporated into many existing machine learning methods. Consequently, we developed a novel algorithm, FIFA, for filtering FFPE artifacts, which uses an EBM model, an interpretable decision-tree-based learning algorithm, to address some of the existing shortcomings. We used four independent cohorts composed of paired lymphoma and cervical cancer samples and a breast cancer cell line with both FF and FFPE samples to define clearly annotated training and test sets and demonstrated improved performance over existing methods. Additionally, FIFA filtering increased relevant biological signals in FFPE breast cancer datasets distinct from the training and testing sets. The EBM framework employed by FIFA is computationally efficient and easily amenable to incorporation of additional datasets due to its generalized additive modeling of features making it straightforward to incorporate new data into existing models dynamically over time.

**Conclusions:** Our novel FFPE variant artifact filtering tool, FIFA, is a marked improvement over existing methods. It can be easily implemented, *post hoc*, to supplement existing somatic calling pipelines, training and inference can be run quickly across most compute environments, and it can be easily updated online as new training data becomes available. Accordingly, FIFA represents an important advance in retrospective cancer genomics research by further enhancing access to the vast stores of FFPE-archived tumor samples currently in existence.

## Background

Formalin fixation and paraffin embedding is a treatment routinely used for long-term conservation of clinical tumor samples. Globally, it is estimated that there are more than 400 million FFPE clinical samples archived [1, 2] including many less common tumor types [3, 4]. Although FFPE samples preserve tissue architecture and cellular morphology, the formalin fixation chemically modifies the sample’s nucleic acids. The process can introduce damage such as nucleic acid fragmentation, DNA crosslinks or strand breaks, and artifactual deamination (which manifests as cytosine-to-thymine mutations in next generation sequencing) that can complicate sequencing [5–7]. Even more challenging, the degree of FFPE-induced degradation varies based on sample age, storage time, and fixation protocol [8] introducing confounding batch effects. Enzymatic DNA repair methods exist but only modestly minimize the prevalence of sequencing artifacts [9–11]. Thus, these samples represent an invaluable yet still underutilized resource for genomic and molecular research [12].

Several computational approaches exist to discriminate FFPE artifacts from real genomic variants. The existing solutions include simple variant allele frequency (VAF) cutoffs [13, 14], ensemble variant calling heuristics [15, 16], or read orientation-based filters such as SOBDetector [17] and FilterByOrientationBias (Mutect2) [18]. Recently, more sophisticated machine learning methods have been developed like FFPolish (logistic regression) [19], Ideafix (random forest and XGBoost) [20], and FFPErase (random forest) [21]. DeepSomatic represents the state-of-the-art deep learning approach employing a convolutional neural network model for FFPE-aware variant calling directly from read alignments [22]. The effectiveness of these approaches is varied and important practical limitations persist. For example, filtering variants using VAF cutoffs may unintentionally exclude genuine subclonal mutations, including clinically actionable variants known to occur at low frequencies [23–26]. While methods like FFPolish and Ideafix incorporate additional features in their predictions, neither considers the broad, genomic context surrounding a variant location. Furthermore, Ideafix requires a Mutect2 formatted variant call format (VCF) file for feature extraction, only makes predictions on deamination events, and uses a predetermined VAF cutoff. As with many deep learning algorithms, DeepSomatic’s pretrained models are difficult to interpret and retraining on private and/or new datasets can be laborious. Furthermore, training custom models with one’s own data or supplementing the pretrained models with additional data likely requires considerable computational resources.

In this study, we first assessed the effectiveness of existing methods using cohorts of diffuse large B-cell lymphoma (DLBCL) samples, both fresh frozen and formalin-fixed, in parallel, therefore, providing a means to robustly identify real and artifactual variants. Through this process, we identified strengths and weaknesses inherent to these methods and, subsequently, designed a novel procedure that minimizes these shortcomings. Our approach for filtering FFPE artifacts (FIFA) has been trained on a large set of 90 paired FF and FFPE samples and uses a feature set that includes characterizations of the local genomic context, i.e. windows around variants, in addition to the properties of reads explicitly overlapping variants commonly used in previous methods. We employ an explainable boosting machine (EBM) framework which provides several benefits. First, EBMs have been shown to have comparable performance to some of the routinely best performing approaches such as XGBoost (https://arxiv.org/html/2410.19098v1; https://arxiv.org/pdf/1909.09223). Second, EBMs are highly interpretable at the global and local level, without the need to estimate computationally expensive attribution metrics such as Shapley values [27]. Accordingly, we provide utility scripts to allow for evaluating feature importance contributions for each variant which can facilitate model evaluation and optimization. Third, because EBMs are, at their core, ensemble decision trees, the training and prediction processes are fast and computationally efficient. Last, because EBMs rely on generalized additive functions to quantify feature contributions for each prediction, models can be easily combined making it straightforward to incorporate new data by training models and simply “averaging” them to previously trained models allowing for the uncomplicated creation of consolidated models, e.g., the *n+1* addition of a new cohort to an existing model or custom cancer-type-specific models. In summary, we developed a new machine learning tool, FIFA, for reducing FFPE artifacts in somatic single nucleotide variant (sSNV) calling and demonstrated its improved performance and practical utility in several contexts. We believe FIFA increases the fidelity of somatic variation calling from FFPE samples and represents an important advance towards realizing the full clinical value of the stores of readily available archival samples.

## Methods

### Analysis of whole genome sequencing and classifying real and artifactual somatic variants

Tumor-Normal whole genome sequencing data was uniformly processed using the standard NYGC somatic single nucleotide variant (sSNV) calling pipeline [28] aligning to reference genome GRCh38. We focused on evaluating performance in high confidence variants, those identified by two or more somatic calling algorithms–Strelka2 [29], Mutect2 [30, 31], or Lancet [32]–since, in practice, the lower confidence calls can be (and commonly are) easily removed as a first-pass, false-positive filter. All specimens used in this study had paired FFPE and FF samples. As such, high confidence variants called in FFPE and identified by at least one caller in FF were classified as real variants.

### Evaluating the landscape of existing tools designed to detect FFPE artifacts

To appraise the performance of several existing methods for filtering artifactual sSNVs calls, we used our collection of variants labeled on the basis of paired FF vs. FFPE contrasting. The simplest filtration procedures apply a variant allele frequency (VAF) cutoff, as FFPE artifacts are expected to occur at low alternative allele frequencies. This simple cutoff can be insufficient in more complicated settings where additional factors can blur any separation between real and artifact variant VAFs, e.g., tumor purity and subclonality, etc. As such, we also used a Z-score cutoff, Z<-0.05, to capture variants with lower VAFs relative to the overall distribution, per sample, independent of their absolute fraction. Additionally, we included two machine learning methods which leverage manually curated features associated with reads supporting an sSNV: 1) Ideafix [20] which relies on an XGBoost model trained on 1.6M deamination variants from 27 paired FF and FFPE exome calls; 2) FFPolish which was trained on Strelka calls [29] made in 2 of the 4 cohorts used here, the HIV+ Tumor Molecular Characterization Project (HTMCP) and the Burkitt Lymphoma Genome Sequencing Project (BLGSP), and learned a 31 feature linear model trained on ∼8.7 M variants. Lastly, we evaluated the performance of a state-of-the-art deep learning (DL) convolutional neural network (CNN) model, DeepSomatic, which makes genome-wide variant calls on read pileups represented as images, building on a previously described method for germline variation calling, DeepVariant [33]. DeepSomatic was run on 10/13 samples because one sample failed while two others had too few real variants which unreasonably skewed the per-sample performance evaluations. Notably, we evaluated performance using both the FF and FFPE DeepVariant models and found better performance with the former, which we, therefore, use throughout this study. All methods evaluated in this comparison were run using default parameters, unless otherwise stated.

### Designing a new tool for filtering FFPE sequencing artifacts (FIFA)

Our aim was to develop a tool that incorporated informative features from prior methods along with newer features with the intent of addressing any shortcomings. In addition, we sought a model that was: 1) easy to train, requiring minimal computational resources, 2) easy to interpret, avoiding blackbox models so individual predictions could be assessed, 3) and easy to update, as access to FFPE fixed samples, particularly those with matched fresh frozen samples, is certain to increase over time, e.g., in newly sequenced cancer types and cohorts. To this end, we developed an explainable boosting machine (EBM) model which satisfies all these properties and has been shown to achieve performance similar to XGBoost in benchmarking comparisons (https://arxiv.org/html/2410.19098v1; https://arxiv.org/pdf/1909.09223). We manually selected a set of 60 features to characterize each variant. This included 23 features used by FFPolish [34] and/or Ideafix [20] which were deemed to be of particular relevance. Fifteen of these features were shown to have high importance in the FFPolish model (absolute logistic regression coefficient >0.25) including VAF, mapping quality, base qualities of mismatches, proximity to the read 3’ end, and fragment length. Twelve of our features were also shared with Ideafix, and all were reported to have area under the curve (AUC) > 0.5 in their models. We supplemented this set with additional features (**Supplementary Table 1B**) describing the window flanking (±500bp) each variant in an effort to explicitly emulate information captured by image-based CNNs like DeepSomatic. To account for inconsistent feature distributions, e.g., due to sample-and cohort-level batch effects, we transformed various numerical features at a per-sample basis using an inverse hyperbolic sine transformation to allow for zero and negative values. Finally, we sought a more robust representation of low frequency artifacts in lieu of absolute or preprocessed VAF. We reasoned that the VAF distribution of FFPE artifacts resembles that of passenger mutations in a cancer context. MOBSTER [35] is a tool designed to distinguish lower frequency neutral passenger mutations from (sub)clonal variants in a cancer setting. We repurposed MOBSTER’s probability of a variant lying in the neutral tail as a proxy for a variant being an FFPE artifact and included it as an additional feature in our model. Using these features, we trained an EBM, using the `interpret (v0.7.3)` python package, on each of our four cohorts independently and combined them in various permutations to create consolidated models and evaluated their performance. In order to demonstrate EBM performance was comparable to state-of-the-art machine learning methods, we also trained a separate XGBoost model, using similar hyperparameterization, for comparison (**Supplementary Tables 1D**).

### Hyperparameter Optimization

We implemented the ExplainableBoostingClassifier (EBM) from the `interpret` package (v0.7.3), which at the time of writing, did not include native functions for hyperparameter optimization. Consequently, we leveraged the open-source framework, Optuna (version 4.5.0) [36], to conduct a stratified 5-fold grid search, optimizing F1 score for five hyperparameters: max_bins, max_leaves, early_stopping_rounds, early_stopping_tolerance, and learning_rate (**Table 1**). We conducted this grid search independently during training for each cohort’s individual model.

**Table 1.** EBM hyperparameters. Optuna was used to search for the optimal hyperparameter sets for each model trained.

| Hyperparameter | Values |
| --- | --- |
| max_bins | { 32, 256, 512, 1024 } |
| max_leaves | { 2, 3, 2005 } |
| early_stopping_rounds | { 50, 100 } |
| early_stopping_tolerance | { 0, 1e-5, 5e-5 } |
| learning_rate | { 0.005, 0.015 } |

### Round-robin testing

As an initial validation, we tested each of our four training cohorts of paired samples (NYGC1, NYGC2, BLGSP, and HTMCP). For each iteration, we held out one of the four cohorts as a test set and trained FIFA by merging the independent models of the remaining three training sets using the `interpret::merge_ebms` function. During our evaluation process, we noted that HTMCP exhibited significant batch effects and produced models that performed poorly (**Supplementary Table 1C**). Therefore, we also explored models excluding HTMCP and carried out the leave-one-cohort-out testing procedure with the remaining three cohorts.

### Testing performance in an independent set of FF and FFPE archived HCC1395 cells

Our final FIFA model–trained on NYGC1, NYGC2, and BLGSP–was applied to the variant callsets generated by Mutect2 for three FFPE-fixed replicates of the HCC1395 cell line previously reported by SEQC2 [37]. The VCFs and binary alignment/map (BAM) files for the replicates were downloaded directly from the SEQC2 consortium and to each we applied FFPolish and a 10% VAF cut-off (https://ftp-trace.ncbi.nlm.nih.gov/ReferenceSamples/seqc/Somatic_Mutation_WG). We compared these results against those produced by DeepSomatic, restricting our analysis to the regions the authors of the tool describe as high confidence. Testing was performed only on chromosome 1 as DeepSomatic used the remaining chromosomes for training purposes.

### Functional assessment of FIFA filtering in the breast cancer cohort, NYGC3

For additional biological validation, we applied FIFA to the high confidence sSNV calls from a cohort of 143 breast cancer FFPE samples (NYGC3) processed by the NYGC somatic variant calling pipeline. We identified 98 samples with matched RNAseq and searched for evidence of expression of the somatic allele to corroborate a variant was real, as the probability of detecting FFPE-induced artifacts at the same base in both DNA and RNA molecules is low. We generated read pileups at the putative variant locations, from alignment BAM files, limiting the scope to coding variants as annotated by the Variant Effect Predictor (VEP). We calculated the coverage at each position as the sum of the number of reads supporting the REF, ALT, or OTHER_ALT alleles in our pileups and evaluated performance at various coverage thresholds. A somatic variant was considered “Real” if there was at least one read supporting the alternate allele. Additionally, we used MuSiCal to decompose the 96-channel single-base substitution mutational signatures present in NYGC3, pre-and post-filtering, with FFPolish and FIFA. We downloaded previously published trinucleotide mutation counts from the International Cancer Genome Consortium (ICGC) [38] and the Genomics England 100,000 Genomes Project (GEL) for two fresh-frozen breast cancer cohorts and, as negative controls, the GEL colorectal cancer and GEL bone and soft tissue cancer cohorts (https://zenodo.org/records/5571551) [39], decomposing mutational signatures from the initial count matrices with MuSiCal, for consistency. Lastly, we predicted homologous recombination deficiency, as associated with BRCA1/BRCA2 mutations, using a machine learning method, HRDetect [40] that leverages mutational signatures, copy number estimates, and structural variation calls. The proportion of the COSMIC single base substitution (SBS) signature 3 was then used to quantify the veracity of the classifications, where higher SBS3 proportion in HRD+ samples suggested improved classification.

## Results

We conducted an evaluation of several common methods for minimizing FFPE artifact sSNVs in B-cell lymphoma samples. In brief, performance was found to be varied with, to our surprise, no clear benefit to more sophisticated methods. In this study, we aimed to develop a novel tool that was fast and easy to use like conventional ML but with performance comparable or surpassing more complex algorithms like the neural networks employed by DeepSomatic. We developed an EBM-based decision tree model for Filtering FFPE Artifacts (FIFA). We used combinations of our 4 cohorts (NYGC1, NYGC2, HTMCP, and BLGSP), composed of paired FF/FFPE samples, to show that FIFA outperformed existing methods in unseen data (leave-one-cohort-out cross-validation). We then formalized a final model using the best combination of cohorts and showed that in a previously published HCC1395 breast cancer cell line dataset [37] and a separate, unpublished cohort of breast cancer tumor samples (NYGC3), FIFA exhibited a notable increase in various biological signals.

### Evaluation of existing methods for reducing FFPE artifactual variant calls

To address the challenge of defining training and evaluation data sets that reliably distinguish between real and artifact variants, prior studies have used paired FF and FFPE samples [41, 42] from the Human Tumour Molecular Characterization Project (HTMCP) [43] and Burkitt Lymphoma Genome Sequencing Project (BLGSP) [44] with the assumption that real variants would be present in both samples while fixation artifacts would be private to the FFPE samples. Building on this concept, we expanded upon previously published pairs with two additional cohorts of 13 DLBCL and 7 B-cell lymphoma FF/FFPE pairs, which we refer to as NYGC1 and NYGC2, respectively, comprising an additional 449,501 high confidence variants. Across all four cohorts, this resulted in 6,087,196 labeled variants (**Table 2** and **Supplementary Table 1A**).

**Table 2.**
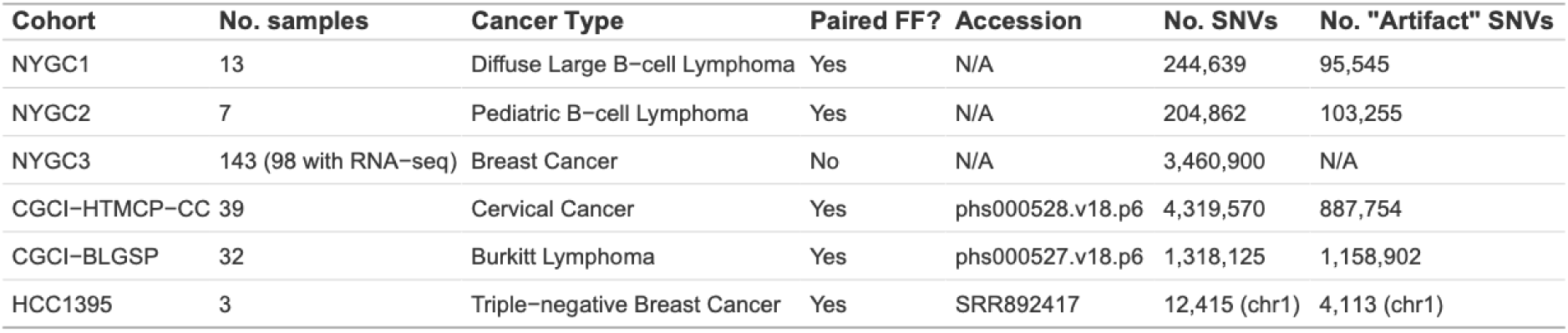
Cohorts used in this study. Four cohorts with paired FF/FFPE samples were used for training and testing (NYGC1, NYGC2, HTMCP, and BLGSP). Two additional FFPE-only cohorts were used exclusively for performance evaluation (HCC1395 and NYGC3). Indicated are the total and artifact number of high-confidence sSNVs called by the NYGC somatic pipeline in each cohort.

We evaluated the performance of simple threshold cutoffs, variant allele frequencies (VAF) less than 5%, or, more permissively, VAF less than 10%. Additionally, we converted VAFs to per-sample Z-scores to account for differences in VAF distributions between samples common in tumor sequencing, e.g., due to varying purity and clonality. We also evaluated the linear model based method, FFPolish [45], and a convolutional neural network, DeepSomatic [46]. Each method was evaluted for its ability to identify artifacts resulting from FFPE-induced damage to DNA in cohorts of 12 diffuse large B-cell lymphoma (DLBCL) samples and 7 B-cell lymphoma samples, which we refer to as NYGC1 and NYGC2, respectively. These data sets were composed of FF and FFPE samples derived from the same tumor. After uniform processing through the New York Genome Center’s (NYGC) somatic variant calling pipeline [47], aligning to reference genome GRCh38, we contrasted sSNV calls from both preservation methods to establish real and artifactual variants. NYGC’s pipeline identifies high confidence calls as those corroborated by multiple calling methods. As an initial filter, we selected for only these high confidence calls, as is common practice, as a first-pass means to reduce false positives from downstream analysis. We focused our performance evaluation on identifying FFPE artifacts in only the remaining calls. FFPE variants supported by any level of calling evidence in FF (low or high confidence) were then labeled as “Real”. All remaining high confidence FFPE variants were labeled as “Artifacts”.

We first evaluated performance in the 13 DLBCL samples in NYGC1. Because Ideafix only filters deamination events after applying a VAF <30% cutoff, we constrained our analysis to C>T deamination mutations below 30% VAF in order to compare tools fairly. We applied Ideafix, FFPolish, and the 5%, 10%, and Z-score VAF cutoffs to the high confidence variants in the uniformly processed FFPE VCFs. For DeepSomatic, which makes genome-wide calls rather than *post hoc* filtering VCFs, we restricted its assessment to only positions that have been labeled as real or artifact mutations, removing from consideration any unlabeled variants that did not intersect high confidence calls from NYGC’s somatic pipeline in FFPE samples. We calculated performance metrics, per sample, and found DeepSomatic had the highest median accuracy (0.89) while FFPolish and Z-score had the highest F1 score (0.83) (**Figure 1A and 1B**, left). We postulated that DeepSomatics’s inclusion of contextual information through image-like representations of the local regions around each evaluated loci may have been of novel value. This genomic context is not captured in most existing, conventional machine learning (ML) methods like FFPolish. Surprisingly, the simple 10% VAF cutoff exhibited higher median accuracy (0.83) and a modest median F1 score (0.74) relative to the more sophisticated ML methods, Ideafix and FFPolish (0.57 and 0.83, respectively). When evaluating the cohort as a whole, aggregating variants from all samples rather than per sample, VAF 10% again performed reasonably well with a 0.74 overall F1 score (**Figure 1C**, left). In summary, performance was varied in discriminating real from artifact deamination events in NYGC1. Unexpectedly, the machine learning methods did not substantively outperform simple VAF cutoffs.

**Figure 1.**
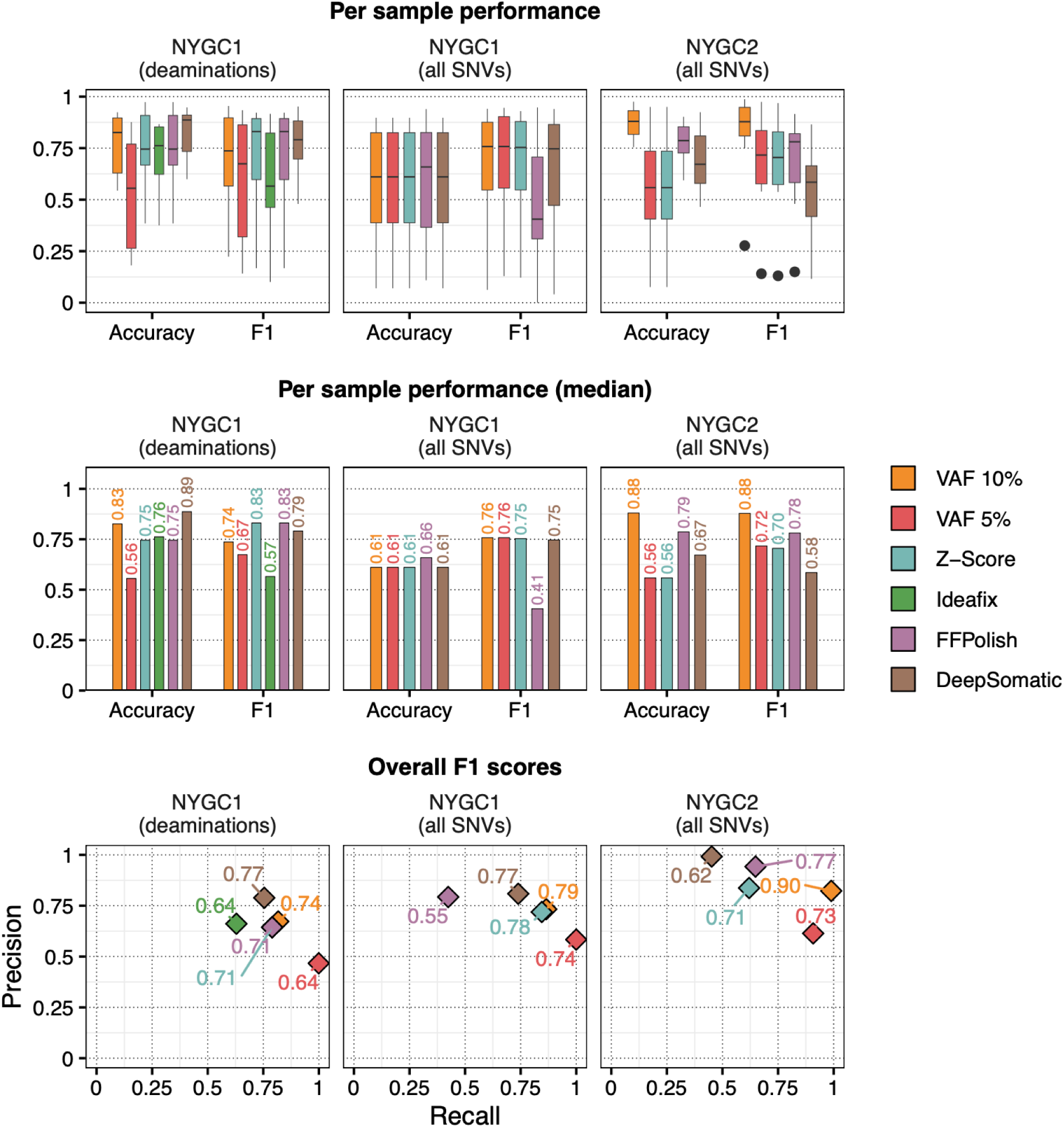
Existing methods showed varied ability to filter FFPE artifacts. **A)** Per sample accuracy and F1 scores for each method in NYGC1 deaminations (left) as well as performance in the full NYGC1 (middle) and NYGC2 (right) call sets. Because Ideafix forces a strict call of “real” at all variants with VAF>30%, we limited the evaluation to only deaminations below this hard threshold. **B)** Median accuracy and F1 scores for each sample for the same three cohorts. **C)** Cohort-level recall, precision, and F1 scores across each of the test sets–all cohort variants were aggregated and evaluated collectively.

Given Ideafix’s limited scope, we decided against including it in subsequent benchmarking. We then expanded our evaluation to include non-deamination variant types across the full VAF spectrum in the NYGC1 cohort. Once again, performance was varied between the tools in terms of accuracy and F1 score. FFPolish displayed the highest accuracy at the expense of reduced recall and F1 score (**Figure 1A-C**, middle). VAF 10% continued to perform comparably to or better than other tools. To get a broader understanding of method performance, we conducted a second evaluation in an independent cohort of 7 paired B-cell lymphoma samples (NYGC2). As before, we limited our analysis to high confidence positions labeled real or artifact mutations. The 10% VAF, yet again, showed surprisingly good performance. Its accuracy and F1 scores, at both the per-sample and cohort-wide level, were substantially higher than all other tools, with the highest per-sample median accuracy (0.88) and F1 (0.88), and cohort-wide, overall F1 (0.90) (**Figure 1C**, right). DeepSomatic’s performance was in line with all other methods in NYGC1 but dropped in NYGC2, particularly its F1 scores. All together, the performance of these tools was, yet again, varied. As in NYGC1, the more complex ML methods failed to consistently outperform the simple 10% VAF cutoff. DeepSomatic’s performance in NYGC1 was contradicted by the drop in sensitivity in NYGC2 speaking to the complexity of this problem where different datasets can experience distinct consequences of FFPE damage making ML methods difficult to generalize. In light of this, we sought to leverage our larger set of four cohorts of paired FF and FFPE samples to advance the current slate of FFPE artifact filtering tools.

### Round robin validation of a novel explainable boosting machine for filtering FFPE artifacts

Our survey of the current landscape of artifact detection methods suggested VAF cutoffs are reasonably effective in limiting FFPE artifacts. However, there remains a concern they will fail in more complex circumstances, i.e. low purity or subclonality, where VAF alone is insufficient to discriminate between real and artifact sSNVs. Nonetheless, the more sophisticated machine learning algorithms performed poorly in our test datasets leaving open the possibility of ineffective incorporation of features, putatively associated with FFPE damage, beyond VAF, into those models. DeepSomatic showed promising results, performing well on NYGC1, but faltered with a drop in F1 score in NYGC2. In principle, it is likely DeepSomatic would exhibit a significant performance improvement if trained on a broader dataset, as we have in this study, rather than only in HCC1395 cell line data. However, in practice, neural networks can be complex and often require extensive computational resources and, thus, have a higher barrier of entry. Therefore, we developed FIFA using a more conventional machine learning method that is easy-to-use and re-train as new data becomes available, is computationally lightweight, given its modest hardware requirements and minimal run time, and, most importantly, performs well.

Ensemble models are well-known for their straightforward implementation and state-of-the art performance [48]. Consequently, we used an explainable boosting machine, an ensemble of generalized additive functions, because their interpretability of individual variant predictions could be leveraged to improve any shortcomings in the model. Additionally, the additive feature functions are simple to “combine”, allowing us to train and combine independent models as new data arrives through a natural process of averaging feature functions together (**Figure 2A**). EBM performance is ostensibly comparable to state-of-the-art methods like XGBoost (https://arxiv.org/html/2410.19098v1; https://arxiv.org/pdf/1909.09223), which was consistent with our experience (**Supplementary Table 1D**).

**Figure 2.**
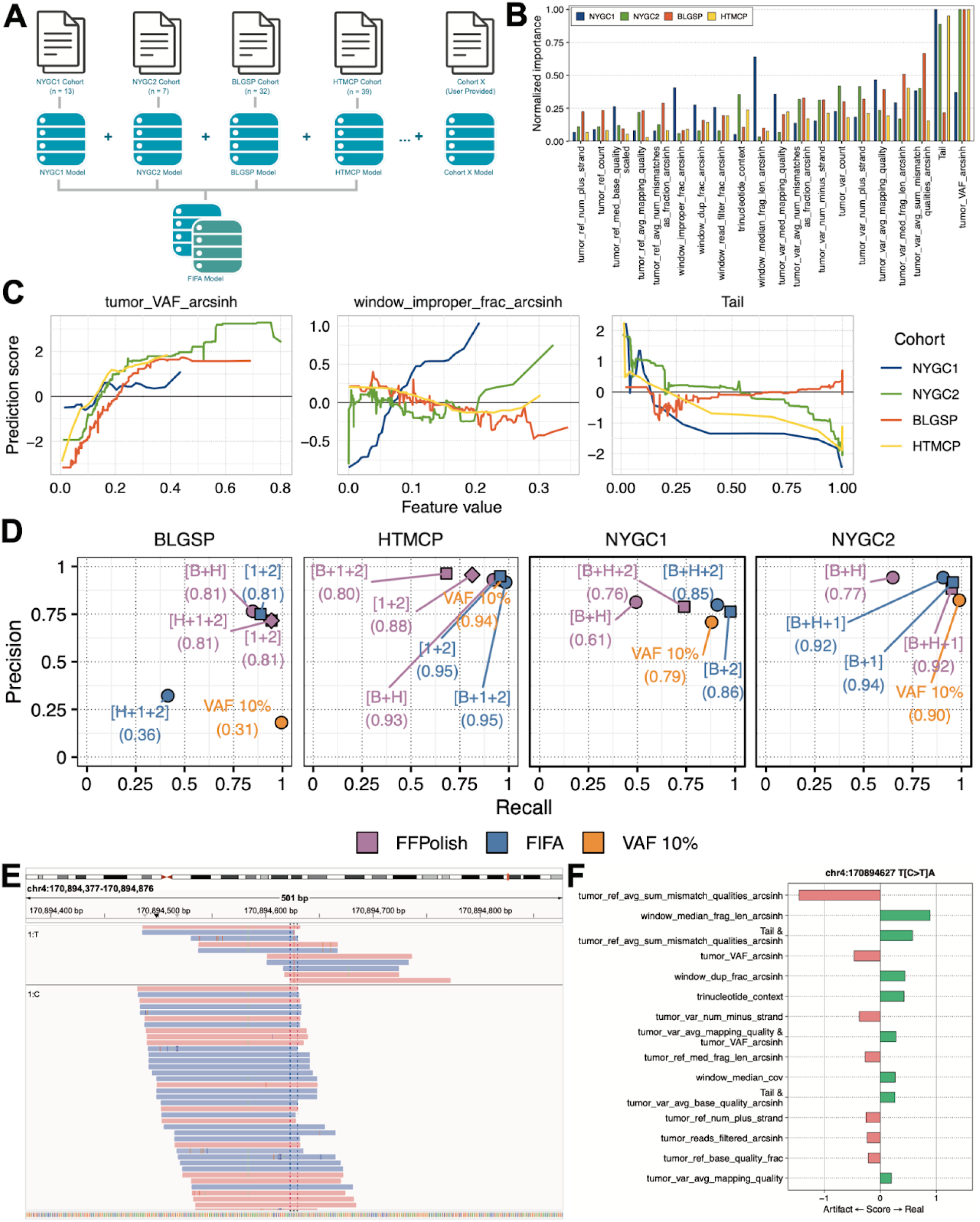
An explainable boosting machine for discriminating real and artifactual somatic SNVs. **A)** Training FIFA requires variant locations and an associated BAM file. We trained models independently for four cohorts and merged and evaluated them in various permutations. This merging process enables updating with the incorporation of new models trained on additional cohorts of data as it becomes available. **B)** Normalized importance for 20 selected features from each of the models trained independently on each of the four cohorts. **C)** Feature score functions for three selected features by cohort model. **D)** Cohort-level performance metrics tested on each of four held out cohorts. Training datasets are indicated as abbreviations within brackets: (B) BLGSP, (H) HTMCP, (1) NYGC1, (2) NYGC2. F1 scores are shown in parentheses. The published FFPolish model was trained in BLGSP and HTMCP. **E)** A selected variant, GRCh38 coordinate chr4:170894627:C>T, identified as real by FIFA but filtered as an artifact by other methods. **F)** Feature contributions for a putative mutation at chr4:170894627:C>,T illustrating how window-based features contributed to the retention of this variant.

We trained EBM models for each of our four cohorts, independently, performing a stratified 5-fold cross-validation to optimize hyperparameters for each. Specifically, we used the Optuna framework of Study objects [36] to efficiently constrain our hyperparameter search space. We extracted feature importances to contrast the differences between each cohort model (**Figure 2B**). While inverse hyperbolic sine transformed tumor VAF was unsurprisingly the most important feature in BLGSP, HTMCP, and NYGC2, several features had higher importance in NYGC1, including the MOBSTER tail probability and several window-based features. e.g., median fragment length, fraction of duplicates, and fraction of improperly paired reads. Inspection of the generalized additive functions for these features illustrated how they are associated differently with “real” predictions across the value range (**Figure 2C**). These differences highlight the need to consider diverse training data and the potential advantage of combining independently trained EBM models in order to generalize more broadly across all possible FFPE-derived data sets.

To gauge how well combining independent models performs, we conducted a round-robin validation, combining the independently pre-trained cohort models and testing on the variants of the fourth held-out cohort. Scoring the variants using a VAF 10% cutoff, FFPolish, and FIFA (combining 2 or 3 models), we found FIFA outperformed the other methods by F1 score when testing in NYGC1, NYGC2, and HTMCP (**Figure 2D**). In BLGSP, the 10% VAF could not distinguish between real and artifact variants because there is substantial overlap between their VAF distributions, and as a result, precision suffered (F1 of 0.20). The published FFPolish model, trained on the BLGSP and HTMCP cohorts, showed poor recall in NYGC1 and NYGC2 (F1 of 0.61 and 0.77, respectively). Notably, FIFA performed poorly in BLGSP (F1 of 0.36), despite high performance in the other three cohorts. After permuting different combinations of independent cohort models, we found that when the HTMCP cohort was used in training, performance was often detrimentally affected (**Supplementary Table 1C**). To proceed, we excluded HTMCP, using only NYGC1 or NYGC2 paired with BLGSP, and performed testing on NYGC2 or NYGC1. We used the same model, trained on NYGC1 and NYGC2, to test in BLGSP and HTMCP. These 2-model versions of FIFA produced the highest F1 scores for each of the cohorts (F1’s of 0.81 to 0.95). To more objectively compare performance, we trained additional FFPolish models using the same permutations of two and three cohorts as with FIFA. FIFA consistently outperformed FFPolish, by F1 score, with the exception of the NYGC1, NYGC2, HTMCP trained model tested in BLGSP. Given this outcome, we combined NYGC1, NYGC2, and BLGSP to produce our final FIFA model. As an illustrative example of this final FIFA model performance, we inspected the region around a high confidence variant call at chr4:170894627, where we observed a deamination that FIFA alone correctly identified as real, due in part to the contribution of three of the novel features we introduced: the median fragment length and duplicate fraction in the window and the tail probability from MOBSTER (as an interaction term [49] with the average reference base mismatch quality scores) (**Figure 2E and 2F**).

### Evaluating FIFA in paired fresh-frozen and FFPE HCC1395 mammary gland epithelial cells

In order to evaluate the generalizability of FIFA in unseen data, we tested its performance in paired FF/FFPE HCC1395 samples, a human breast cancer cell line dataset reported previously by the SEQC2 consortium [37]. We evaluated the performance of FIFA against DeepSomatic, VAF 10%, and an XGBoost model trained with similar hyperparameters to our EBM. It should be noted that DeepSomatic was modeled on this cell line data–holding out chromosome 1 for testing and training on the remaining chromosomes. In light of this, we restricted our evaluation to only the chr1 “test” variants. We used the preprocessed Mutect2 sSNV calls produced by SEQC2 and applied *post-hoc* filtering with FIFA, VAF10%, and our XGBoost model and compared predictions with the reported DeepSomatic results (**Figure 3**). By applying FIFA to the Mutect2 calls, we dramatically improved recall, recovering nearly every real variant, with a small reduction in precision. Across the replicates, FIFA outperformed other variant filters with a mean F1 score of 0.961 compared to 0.880-0.933 for the other methods (**Supplementary Table 1C**). We recognize that the intended use case for DeepSomatic is as an end-to-end somatic variant calling pipeline. *Post-hoc* filtering of Mutect2 calls with FIFA outperforms unfiltered DeepSomatic calls. However, all filtering methods only marginally improve DeepSomatic’s sSNV calls, highlighting the strong overall performance of the method as a somatic SNV caller. By supplementing with FIFA filtering, a simple, conventional Mutect2 sSNV calling pipeline can obtain similar or improved performance compared to state-of-the-art DL methods like DeepSomatic.

**Figure 3.**
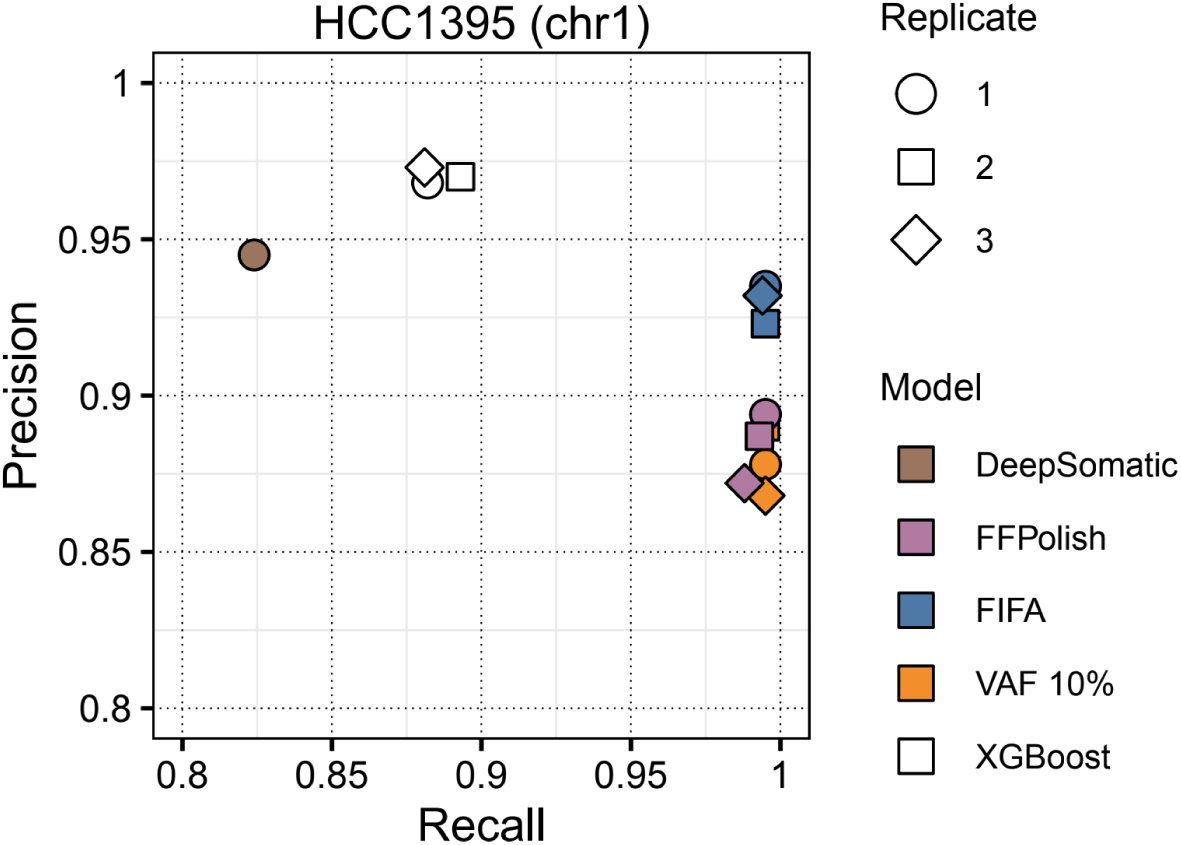
FIFA outperforms other methods in a human breast cancer cell line, HCC1395. Precision and recall metrics for various methods in the HCC1395 dataset composed of 3 separate replicates previously published by SEQC2. Performance is shown for 1) FFPolish, both the published, original model and a version retrained on the same data as FIFA, 2) FIFA, 3) VAF 10% cutoff, 4) XGBoost parameterized similarly to FIFA, 5) DeepSomatic, showing only the aggregated metrics as reported previously in their manuscript.

### Functional corroboration of FIFA predictions in a breast cancer cohort

We leveraged a cohort 143 FFPE-preserved, breast cancer tumor-normal WGS samples (NYGC3), with matched bulk tumor RNA-seq data to further validate FFPE artifact filtering performance. With no paired FF samples to corroborate the veracity of called sSNV mutations we first searched for reads supporting the somatic allele in the RNA-seq data as an alternative functional corroboration of real variants. We reasoned that FFPE artifacts were unlikely to co-occur at identical base positions in both DNA and RNA molecules and used the paired RNA-seq data to confirm expression of the somatic allele as corroboration of a variant’s authenticity. We evaluated the frequency with which the alternative allele could be observed in bulk RNA-seq derived from the same sample as corroboration of a real variant. RNA-seq read coverage over candidate exonic sSNVs was variable with as many as 21,326 variants covered at 5X or greater and 10,785 at 50X or greater (**Figure 4B**). We performed our evaluation at the 12,898 variants with at least 30X coverage where there would be a 95% chance of observing one or more reads supporting a variant with a 10% VAF. FIFA outperformed FFPolish and 10% VAF cutoffs with a noticeably higher precision (**Figure 4A**). Unsurprisingly, performance metrics degraded at lower coverages, particularly for low VAF variants, which can be especially difficult since both low probability sampling or lack of expression could explain the absence of read support for a given variant. Nevertheless, FIFA’s predictions showed better corroboration by the matched RNA-seq data compared to other methods.

**Figure 4.**
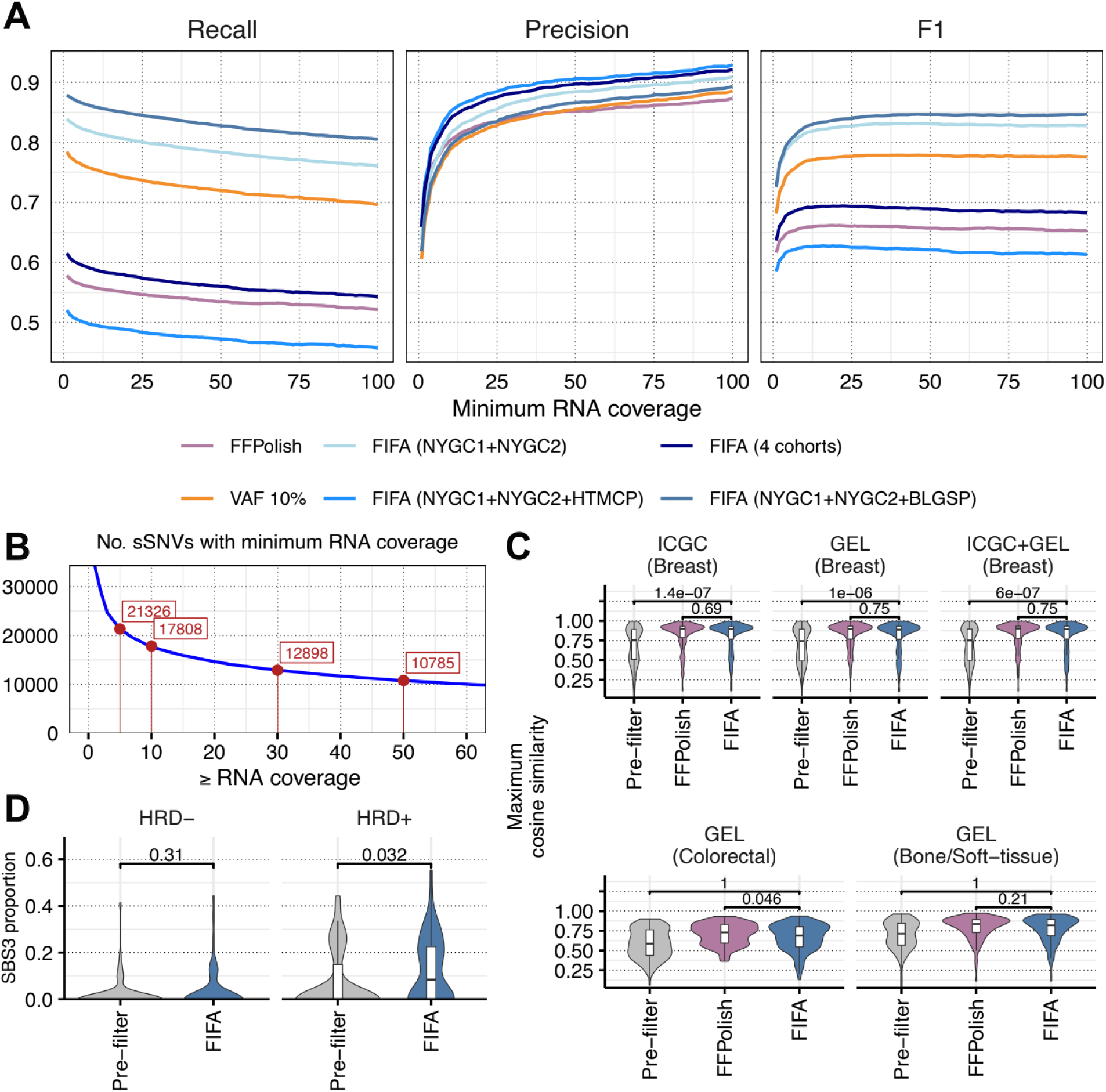
Functional validation of sSNVs in breast cancer cohort NYGC3. **A)** We identified all putative exonic sSNVs in our breast cancer cohort, NYGC3, covered by matched bulk RNA-seq at various read depths from the same tumor and used reads carrying the alternative allele as evidence the variant was real. **B)** Mutation counts were aggregated per sample and deconvolved into compositional mutational signatures. The maximum cosine similarity between each NYGC3 sample and samples from the four public cohorts, ICGC and GEL breast and GEL colorectal and bone/soft-tissue, was determined before and after filtering by either FFPolish or FIFA. **C)** The maximum cosine similarity of mutational signature proportions was calculated between NYGC and the ICGC and GEL cohorts. Filtering methods like FFPolish and FIFA substantially improved the best match suggesting an enrichment for mutations observed in external fresh frozen cohorts. **D)** The mutational signature, SBS3, is associated with homologous recombination deficiencies (HRD) and BRCA1 and BRCA2.

As an additional functional corroboration, we surmised that successful filtering of FFPE artifacts should retain variants representative of expected mutational signatures. To determine whether this was in fact the case, we decomposed mutation counts for samples from the NYGC3, ICGC, and GEL cohorts into contributions of known COSMIC mutational signatures (v3.4, GRCh37, https://cancer.sanger.ac.uk/signatures/sbs/) [50, 51] using MuSiCal [52]. Lacking truth information, we reasoned that our cohort would likely reflect the mutation spectrum of other previously published breast cancer samples. Consequently, for each of our samples, we identified the maximum cosine similar sample from either ICGC (n=741) or GEL (n=2572) breast cancer cohorts, and hypothesized that, overall, the maximum cosine similarities for each sample should increase significantly, post-filtering, if artifactual variants were being removed selectively. We found that decomposition proportions were significantly more similar to the ICGC and/or GEL cohorts with filtering than without (**Figure 4C**). Conversely, we performed a similar comparison to unrelated GEL cancer cohorts, colorectal (n=2348) and bone/soft-tissue (n=1480). Artifact filtering did modestly increase the maximum cosine similarity, but less so in FIFA compared to FFPolish (significantly less in colorectal cancer). From this, we concluded that FFPE filtering may be crucially important prior to mutational signature decomposition. While FFPolish and FIFA were both effective at enriching cancer signatures, FIFA was more specific, showing less similarity to unrelated colorectal and bond/soft-tissue cancers.

For one final validation, we tested whether signature decomposition, specifically of SBS3–known to be associated with mutations in BRCA1/BRCA2, and homologous recombination deficiency (HRD) [53]–was correlated with HRD status. To do so, we utilized HRDetect [54] to classify patients with or without HRD and compared their SBS3 proportions, before and after FFPE artifact filtering. We measured the SBS3 proportion inferred by MuSiCal in samples identified as HRD+ versus HRD-by HRDetect, comparing with and without FIFA filtering, and found that FIFA significantly increased the SBS3 proportion in HRD+ samples (**Figure 4D**), providing further evidence of enrichment of relevant biological signal through FFPE-artifact filtering.

## Discussion

In this study, we evaluated a collection of representative methods for minimizing FFPE artifactual sSNVs caused by FFPE damage and found performance highly variable across samples and cohorts revealing the need for more robust, reliable methods. As a result, we developed a novel machine learning method, FIFA, that uses an explainable boosting machine, a relatively recent innovation in ensemble learning, to address some of the remaining shortcomings. In brief, FIFA supplements previously used features with additional characterizations aimed at capturing local context around variant calls and uses a unique representation of VAF, i.e. the MOBSTER Tail probability, that serves as an alternative means to describe low frequency somatic variants. FIFA shows promising performance in our evaluations. As outlined here, FIFA showed top-of-the-line F1 scores in all four cohorts evaluated during our round-robin leave-one-cohort-out testing. Furthermore, in unseen test datasets, such as the HCC1395 breast cancer cell line and our additional breast cohort, NYGC3, FIFA demonstrated high quality filtering and maintained the increase in similarity to known breast cancer mutational signatures while minimizing the similarity to off-target non-breast cancer samples. The deconvolved signatures, after FIFA filtering, showed a significant increase in SBS3 proportions in samples exhibiting evidence of HRD. FIFA is straightforward to run on most personal computers. We evaluated its execution metrics by training and testing on 100,000 variants on the NYGC compute cluster which it completed in approximately 70 minutes, largely feature extraction, using 4 CPUs and less than 2.75G RAM (**Supplementary Table 1E**). FIFA is agnostic to the processing pipeline, requiring any VCF and its associated BAM as input. Because FIFA’s EBM uses simple generalized additive functions to infer a feature’s contribution to a prediction, models can be easily averaged together making it straightforward to combine independent models or update existing models with new data. One limitation of our study, as with many similar efforts, was the limited training data–using only four cohorts of lymphoma and cervical cancer for training. We would expect this limited training to constrain performance but FIFA’s ability to easily incorporate new data would be especially practical. Accordingly, we plan to update our model as new paired FF/FFPE data becomes available. It should also be noted, FIFA expects high coverage tumor-normal whole-genome sequencing; we have not evaluated its performance in targeted, exome, or tumor-only sequencing nor have we evaluated performance in lower coverage studies. Additionally, while other recent methods have addressed small insertion and deletion artifacts and structural variation [21], we have not but hope to extend our process to tackle these artifacts in the near future.

In conclusion, we present FIFA as a democratic, pipeline-agnostic solution to reducing FFPE artifactual contamination in sSNV call sets. The tool is freely available for academic use at www.github.com/nygenome/fifa. Modules for training, predicting, and appending new data are provided. Additional utility scripts for inspecting individual predictions, examining generalized additive feature functions, and for incorporating paired RNA-seq data (BAMs) for corroborating predictions are also available.

## Conclusions

Enabling accurate and dependable sequencing from the vast store of FFPE archived samples has been a long held ambition and would greatly buttress efforts for retrospective analysis to spur on new findings and direction to discover new clinical treatments. Doing so requires precise and sensitive pipelines able to sift through the vast raft of artifactual variants often found in FFPE-archived tumor samples. We present a novel machine learning tool, FIFA, that is easily implemented, requiring no special hardwares like GPUs or large memory computers. The outputs are interpretable allowing for close manual inspection. The process is amendable, allowing for addition of new models, as new data becomes available, without having to retrain the entire corpus.

## List of abbreviations

FFPE: Formalin-fixed paraffin-embedded
FF: Fresh frozen
EBM: Explainable boosting machine
VAF: Variant allele frequency
VCF: Variant call format
DLBCL: Diffuse large B-cell lymphoma
FIFA: Filtering FFPE artifacts
sSNV: Somatic single nucleotide variant CNN: Convolutional neural network
HTMCP: HIV+ Tumor Molecular Characterization Project
BLGSP: Burkitt Lymphoma Genome Sequencing Project
AUC: Area under the curv
BAM: Binary alignment/map
ICGC: International Cancer Genome Consortium
GEL: Genomics England 100,000 Genomes Project
HRD: Homologous recombination deficiency
CPU: Central processing unit
RAM: Random access memory

## Declarations

V.G.’s work was supported by the MacMillan Family Foundation as part of the MacMillan Center for the Study of the Non-Coding Cancer Genome at the New York Genome Center. The NYGC3 cohort was funded in part by NYGC’s Polyethnic-1000 initiative and a Weill-Cornell EIPM/Illumina partnership, and is being analyzed in the Cancer Grand Challenge SAMBAI project. The method described herein is the subject of a pending U.S. patent application (App. No. 63/998,460) filed by [New York Genome Center/ROBINE].

## References

1. Sah S, Chen L, Houghton J, Kemppainen J, Marko AC, Zeigler R, et al. Functional DNA quantification guides accurate next-generation sequencing mutation detection in formalin-fixed, paraffin-embedded tumor biopsies. Genome Medicine. 2013;5:1–12.

2. Wei L, Dugas M, Sandmann S. SimFFPE and FilterFFPE: improving structural variant calling in FFPE samples. GigaScience. 2021;10:giab065.

3. Robbe P, Popitsch N, Knight SJL, Antoniou P, Becq J, He M, et al. Clinical whole-genome sequencing from routine formalin-fixed, paraffin-embedded specimens: pilot study for the 100,000 Genomes Project. Genetics in medicine: official journal of the American College of Medical Genetics. 2018;20.

4. Sosinsky A, Ambrose J, Cross W, Turnbull C, Henderson S, Jones L, et al. Insights for precision oncology from the integration of genomic and clinical data of 13,880 tumors from the 100,000 Genomes Cancer Programme. Nat Med. 2024;30:279–89.

5. Srinivasan M, Sedmak D, Jewell S. Effect of fixatives and tissue processing on the content and integrity of nucleic acids. Am J Pathol. 2002;161:1961–71.

6. de Schaetzen van Brienen L, Larmuseau M, Van der Eecken K, De Ryck F, Robbe P, Schuh A, et al. Comparative analysis of somatic variant calling on matched FF and FFPE WGS samples. BMC Med Genomics. 2020;13:94.

7. Haile S, Corbett RD, Bilobram S, Bye MH, Kirk H, Pandoh P, et al. Sources of erroneous sequences and artifact chimeric reads in next generation sequencing of genomic DNA from formalin-fixed paraffin-embedded samples. Nucleic acids research. 2019;47.

8. Bhagwate AV, Liu Y, Winham SJ, McDonough SJ, Stallings-Mann ML, Heinzen EP, et al. Bioinformatics and DNA-extraction strategies to reliably detect genetic variants from FFPE breast tissue samples. BMC Genomics. 2019;20:1–10.

9. Mathieson W, Thomas GA. Why Formalin-fixed, Paraffin-embedded Biospecimens Must Be Used in Genomic Medicine: An Evidence-based Review and Conclusion. Journal of Histochemistry and Cytochemistry. 2020;68:543.

10. Steiert TA, Parra G, Gut M, Arnold N, Trotta JR, Tonda R, et al. A critical spotlight on the paradigms of FFPE-DNA sequencing. Nucleic acids research. 2023;51.

11. Munchel S, Hoang Y, Zhao Y, Cottrell J, Klotzle B, Godwin AK, et al. Targeted or whole genome sequencing of formalin fixed tissue samples: potential applications in cancer genomics. Oncotarget. 2015;6:25943–61.

12. Guo P, Chen Y, Mao L, Cardilla A, Lee CN, Cui Y, et al. Spatial profiling of chromatin accessibility in formalin-fixed paraffin-embedded tissues. Nature Communications. 2025;16:1–11.

13. Guo Q, Lakatos E, Bakir IA, Curtius K, Graham TA, Mustonen V. The mutational signatures of formalin fixation on the human genome. Nature Communications. 2022;13:1–14.

14. Basyuni S, Heskin L, Degasperi A, Black D, Koh GCC, Chmelova L, et al. Large-scale analysis of whole genome sequencing data from formalin-fixed paraffin-embedded cancer specimens demonstrates preservation of clinical utility. Nature Communications. 2024;15:1–12.

15. de Schaetzen van Brienen L, Larmuseau M, Van der Eecken K, De Ryck F, Robbe P, Schuh A, et al. Comparative analysis of somatic variant calling on matched FF and FFPE WGS samples. BMC Med Genomics. 2020;13:94.

16. Srinivasan M, Sedmak D, Jewell S. Effect of fixatives and tissue processing on the content and integrity of nucleic acids. Am J Pathol. 2002;161:1961–71.

17. Diossy M, Sztupinszki Z, Krzystanek M, Borcsok J, Eklund AC, Csabai I, et al. Strand Orientation Bias Detector to determine the probability of FFPE sequencing artifacts. Brief Bioinform. 2021;22:bbab186.

18. Cibulskis K, Lawrence MS, Carter SL, Sivachenko A, Jaffe D, Sougnez C, et al. Sensitive detection of somatic point mutations in impure and heterogeneous cancer samples. Nat Biotechnol. 2013;31:213–9.

19. Dodani DD, Nguyen MH, Morin RD, Marra MA, Corbett RD. Combinatorial and Machine Learning Approaches for Improved Somatic Variant Calling From Formalin-Fixed Paraffin-Embedded Genome Sequence Data. Frontiers in genetics. 2022;13.

20. Tellaetxe-Abete M, Calvo B, Lawrie C. Ideafix: a decision tree-based method for the refinement of variants in FFPE DNA sequencing data. NAR Genom Bioinform. 2021;3:lqab092.

21. Domenico D, Gundem G, Levine MF, Arango-Ossa JE, Robbe P, Asimomitis G, et al. Enabling whole genome sequencing analysis from FFPE specimens in clinical oncology. Nat Commun. 2025;16:10649.

22. Park J, Cook DE, Chang P-C, Kolesnikov A, Brambrink L, Mier JC, et al. Accurate somatic small variant discovery for multiple sequencing technologies with DeepSomatic. Nat Biotechnol. 2025. 10.1038/s41587-025-02839-x.

23. Bhagwate AV, Liu Y, Winham SJ, McDonough SJ, Stallings-Mann ML, Heinzen EP, et al. Bioinformatics and DNA-extraction strategies to reliably detect genetic variants from FFPE breast tissue samples. BMC Genomics. 2019;20:1–10.

24. Alborelli I, Bratic HI, Chijioke O, Prince SS, Bubendorf L, Leuenberger LP, et al. Robust assessment of tumor mutational burden in cytological specimens from lung cancer patients. Lung cancer (Amsterdam, Netherlands). 2020;149.

25. Basyuni S, Heskin L, Degasperi A, Black D, Koh GCC, Chmelova L, et al. Large-scale analysis of whole genome sequencing data from formalin-fixed paraffin-embedded cancer specimens demonstrates preservation of clinical utility. Nature Communications. 2024;15:1–12.

26. Robbe P, Popitsch N, Knight SJL, Antoniou P, Becq J, He M, et al. Clinical whole-genome sequencing from routine formalin-fixed, paraffin-embedded specimens: pilot study for the 100,000 Genomes Project. Genetics in medicine: official journal of the American College of Medical Genetics. 2018;20.

27. Watson DS. Interpretable machine learning for genomics. Hum Genet. 2022;141:1499–513.

28. Arora K, Shah M, Johnson M, Sanghvi R, Shelton J, Nagulapalli K, et al. Deep whole-genome sequencing of 3 cancer cell lines on 2 sequencing platforms. Scientific Reports. 2019;9:1–13.

29. Kim S, Scheffler K, Halpern AL, Bekritsky MA, Noh E, Källberg M, et al. Strelka2: fast and accurate calling of germline and somatic variants. Nat Methods. 2018;15:591–4.

30. Benjamin D, Sato T, Cibulskis K, Getz G, Stewart C, Lichtenstein L. Calling Somatic SNVs and Indels with Mutect2. bioRxiv. 2019.

31. McKenna A, Hanna M, Banks E, Sivachenko A, Cibulskis K, Kernytsky A, et al. The Genome Analysis Toolkit: a MapReduce framework for analyzing next-generation DNA sequencing data. Genome Res. 2010;20:1297–303.

32. Narzisi G, Corvelo A, Arora K, Bergmann EA, Shah M, Musunuri R, et al. Genome-wide somatic variant calling using localized colored de Bruijn graphs. Commun Biol. 2018;1:20.

33. Poplin R, Chang P-C, Alexander D, Schwartz S, Colthurst T, Ku A, et al. A universal SNP and small-indel variant caller using deep neural networks. Nat Biotechnol. 2018;36:983–7.

34. Dodani DD, Nguyen MH, Morin RD, Marra MA, Corbett RD. Combinatorial and Machine Learning Approaches for Improved Somatic Variant Calling From Formalin-Fixed Paraffin-Embedded Genome Sequence Data. Frontiers in genetics. 2022;13.

35. Caravagna G, Heide T, Williams MJ, Zapata L, Nichol D, Chkhaidze K, et al. Subclonal reconstruction of tumors by using machine learning and population genetics. Nat Genet. 2020;52:898–907.

36. Akiba T, Sano S, Yanase T, Ohta T, Koyama M. Optuna: A next-generation hyperparameter optimization framework. arXiv [cs.LG]. 2019.

37. Fang LT, Zhu B, Zhao Y, Chen W, Yang Z, Kerrigan L, et al. Establishing community reference samples, data and call sets for benchmarking cancer mutation detection using whole-genome sequencing. Nat Biotechnol. 2021;39:1151–60.

38. Zhang J, Bajari R, Andric D, Gerthoffert F, Lepsa A, Nahal-Bose H, et al. The International Cancer Genome Consortium Data Portal. Nat Biotechnol. 2019;37:367–9.

39. Degasperi A, Zou X, Amarante TD, Martinez-Martinez A, Koh GCC, Dias JML, et al. Substitution mutational signatures in whole-genome-sequenced cancers in the UK population. Science. 2022;376.

40. Davies H, Glodzik D, Morganella S, Yates LR, Staaf J, Zou X, et al. HRDetect is a predictor of BRCA1 and BRCA2 deficiency based on mutational signatures. Nat Med. 2017;23:517–25.

41. de Schaetzen van Brienen L, Larmuseau M, Van der Eecken K, De Ryck F, Robbe P, Schuh A, et al. Comparative analysis of somatic variant calling on matched FF and FFPE WGS samples. BMC Med Genomics. 2020;13:94.

42. Dodani DD, Nguyen MH, Morin RD, Marra MA, Corbett RD. Combinatorial and Machine Learning Approaches for Improved Somatic Variant Calling From Formalin-Fixed Paraffin-Embedded Genome Sequence Data. Frontiers in genetics. 2022;13.

43. Gagliardi A, Porter VL, Zong Z, Bowlby R, Titmuss E, Namirembe C, et al. Analysis of Ugandan cervical carcinomas identifies human papillomavirus clade-specific epigenome and transcriptome landscapes. Nat Genet. 2020;52:800–10.

44. Grande BM, Gerhard DS, Jiang A, Griner NB, Abramson JS, Alexander TB, et al. Genome-wide discovery of somatic coding and noncoding mutations in pediatric endemic and sporadic Burkitt lymphoma. Blood. 2019;133:1313–24.

45. Dodani DD, Nguyen MH, Morin RD, Marra MA, Corbett RD. Combinatorial and Machine Learning Approaches for Improved Somatic Variant Calling From Formalin-Fixed Paraffin-Embedded Genome Sequence Data. Frontiers in genetics. 2022;13.

46. Park J, Cook DE, Chang P-C, Kolesnikov A, Brambrink L, Mier JC, et al. Accurate somatic small variant discovery for multiple sequencing technologies with DeepSomatic. Nat Biotechnol. 2025. 10.1038/s41587-025-02839-x.

47. Arora K, Shah M, Johnson M, Sanghvi R, Shelton J, Nagulapalli K, et al. Deep whole-genome sequencing of 3 cancer cell lines on 2 sequencing platforms. Scientific Reports. 2019;9:1–13.

48. Mohammed A, Kora R. A comprehensive review on ensemble deep learning: Opportunities and challenges. J King Saud Univ - Comput Inf Sci. 2023;35:757–74.

49. Accurate Intelligible Models with Pairwise Interactions. https://www.cs.cornell.edu/~yinlou/papers/lou-kdd13.pdf. Accessed 25 Nov 2025.

50. Sondka Z, Dhir NB, Carvalho-Silva D, Jupe S, Madhumita, McLaren K, et al. COSMIC: a curated database of somatic variants and clinical data for cancer. Nucleic Acids Res. 2024;52:D1210–7.

51. Alexandrov LB, Kim J, Haradhvala NJ, Huang MN, Tian Ng AW, Wu Y, et al. The repertoire of mutational signatures in human cancer. Nature. 2020;578:94–101.

52. Jin H, Gulhan DC, Geiger B, Ben-Isvy D, Geng D, Ljungström V, et al. Accurate and sensitive mutational signature analysis with MuSiCal. Nat Genet. 2024;56:541–52.

53. Nik-Zainal S, Davies H, Staaf J, Ramakrishna M, Glodzik D, Zou X, et al. Landscape of somatic mutations in 560 breast cancer whole-genome sequences. Nature. 2016;534:47–54.

54. Davies H, Glodzik D, Morganella S, Yates LR, Staaf J, Zou X, et al. HRDetect is a predictor of BRCA1 and BRCA2 deficiency based on mutational signatures. Nat Med. 2017;23:517–25.

